# T cell interaction partners of DHHC20

**DOI:** 10.1101/2024.02.14.580290

**Authors:** Deana Haxhiraj, Benno Kuropka, Eva Brencher, David Rosenberger, Helena Brandt, David Speck, Patrick Scheerer, Britta Brügger, Cecilia Clementi, Christian Freund

## Abstract

Reversible palmitoylation of proteins at cysteine residues represents a post-translational modification that can alter the cellular localization of proteins, change their distribution within lipid membranes or modulate their conformation and molecular interaction patterns. DHHC enzymes catalyze protein palmitoylation, while thioesterases or hydrolases can rapidly remove the acyl-chain. In human T cells, DHHC proteins have been shown to modify proteins that are involved in major signaling pathways such as Ca^2+^-signaling or kinase-dependent activation of transcription factors for cytokines. For DHHC20, a role in the palmitoylation of the Orai1 Ca^2+^-channel has been demonstrated, but otherwise its T cell interaction partners are largely unknown. Here, we show that recombinantly expressed DHHC20 robustly interacts with 28 proteins from Jurkat T cells, as shown by affinity enrichment combined with mass spectrometric analysis. We find a robust interaction between DHHC20 and the β-subunit of the trimeric G protein Gαsβ1γ2, while the typically palmitoylated Ga subunit is not identified. Cross-linking mass spectrometry with purified DHHC20 and Gαsβ1γ2 then confirms a direct interaction between the β1γ2 domains and the enzyme, while rigid docking offers structural poses that are in agreement with the observed intermolecular cross-linking constraints. Thus, we suggest a model where the β1γ2 subunits of a trimeric G protein serve as a stable interaction partner of a DHHC enzyme, presumably acting as a landing platform for the Gα subunit that is subsequently palmitoylated by the enzyme.

## Introduction

Palmitoylation of proteins is a critical post-translational modification that changes the transport properties, anchorage at or partitioning within biological membranes and the conformation of the acylated substrate^1^. The cellular consequences of the altered molecular properties of palmitoylated proteins can then affect intracellular signal transmission in several ways, for example by affecting receptor signaling^2,3^, kinase function^4,5^ or the translocation of transcription factors^6^. A class of enzymes, the so-called DHHC protein acyltransferases (DHHC-PATs) named after their active site residues, is responsible for the reversible formation of a thioester between the palmitic acid and the cysteine of the target protein^7,8^. A two-step mechanism has been proposed to govern the catalytic mechanism, with auto-palmitoylation of the DHHC protein at its active site as a first step and subsequent transfer of the palmitoyl-group to a substrate protein by trans-esterification^9^.

Of the 23 human DHHC members, only a few are structurally characterized, revealing the positioning of the catalytic Zn finger domain with regard to the transmembrane domain and the localization of the acyl chain of the irreversible inhibitor 2-bromo-palmitate^10^. While some sequence motifs are also conserved in the adjacent N- and C-terminal regions of the transmembrane-Zn finger unit, there is also a large degree of uncertainty in the AlphaFold predictions of these parts of the protein^11,12^. Within the target proteins, no clear consensus motifs for the palmitoylated cysteines can be derived from sequence comparisons. Membrane proximity of the modifiable residue is often a good indication in case that transmembrane proteins are acylated, but even then, it is not clear which additional features contribute to substrate recognition and specificity. Moreover, in some cases DHHC enzymes require the chaperoning of other proteins to function as stable and active palmitoyl transferases. The best described example is the stabilization of DHHC9 by the Golgin subfamily member GCP16^13^, with a recent complex structure showing the mode of interaction^14^. The requisite interaction facilitating HRAS and NRAS palmitoylation has been established, while similar complexes involving GCP16 have been demonstrated to be required for palmitoylation by DHHC14 and DHHC18^14^. Thus, these interacting molecules might well modulate substrate specificity upon complex formation.

In T cells, a number of DHHC enzymes have been characterized, in some cases at the proteomic level only^15,16^, in other cases by clearly assigned functionality^5,17^. For example, the localization of Orai1 channels was shown to depend on palmitoylation at Cys-143 by DHHC20 and impaired recruitment of the C143A mutant to the immunological synapse was observed^17^. The C143A mutant Orai1 displays diminished calcium signals, decreased NFAT activation and diminished IL-2 production, accompanied by a reduction in the number of T cell receptors delivered to the synapse. In another study, DHHC21 was described as a Ca^2+^/calmodulin-dependent enzyme that is critical for activation of naïve CD4+ T cells in response to T cell receptor stimulation^18^. Thus, a prominent role of DHHC enzymes in T cell physiology arises and poses the challenge to identify other interaction partners and substrates of PATs in T cells.

Towards this goal we performed pulldown experiments with purified DHHC20 protein to identify interaction partners of DHHC20 derived from T cells. Furthermore, we described a new approach to obtain initial molecular information on DHHC-substrate pairs, combining a conventional pulldown approach with subsequent cross-linking mass spectrometry of purified proteins. In the conventional pulldown experiment conducted herein, 28 possible candidates were confirmed in at least 2 of 3 experiments by LC-MS, some of which represent known palmitoylated proteins. Focusing on the trimeric G protein Gαsβ1γ2 we then perform two XL-MS experiments with purified DHHC20 protein that reveal numerous intra-as well as a few interprotein cross-links. Interestingly, we identify the βγ subunit of the G protein as the interaction partner of DHHC20, indicating that the heterodimer serves as the landing platform for the enzyme, allowing Gαs to be palmitoylated once the trimeric G protein assembles.

## Results

### Expression and characterization of DHHC20

A mammalian expression system in HEK293-6 EBNA was used to express and purify DHHC20 protein tagged on the C-terminus with a dual Strep-Tag to perform biochemical and biophysical experiments. After the initial lysis step, employing 0.1% n-dodecyl-β-D-maltoside (DDM), the soluble proteins were separated from the membrane fraction through centrifugation. Subsequently, the membrane and organelle fraction containing DHHC20 was lysed utilizing 1% DDM for efficient solubilization. This two-step strategy proved instrumental in achieving a pure preparation of DHHC20, as the second lysis originated from a more refined and purified membrane fraction. The purified DHHC20 was separated by SDS-PAGE as shown in **Fig. S1a**, with DHHC20 being the predominant component. Additionally, traces of the chaperone HSP70 were observed, likely binding to a minor fraction of DHHC20, which can be attributed to its overexpression. The integrity of the protein was confirmed by CD spectroscopy (**Fig. S1b**).

### Interaction partners of DHHC20 in Jurkat T cells

Subsequently, we utilized the bead-bound DHHC20 protein as a bait to perform pulldown experiments in order to identify interaction partners from Jurkat T cell lysates as illustrated in the workflow depicted in **Fig. 1a**. Three biological replicates of the pulldown experiment were performed. As a control experiment, empty beads were incubated with Jurkat T cell lysate to distinguish proteins enriched by DHHC20 from background proteins that bind to the beads. The Strep-tagged DHHC20 and bound proteins were eluted from the beads using biotin and all proteins were identified and quantified by liquid chromatography-mass spectrometry (LC-MS) using label-free quantification (**Table S1**). All proteins with at least a 4-fold increase in relative intensity compared to control (log2 fold difference >2) and a q-value <0.05 were considered significantly enriched. Looking exemplarily at pulldown experiment 1, a total number of 2180 proteins was quantified, and 78 proteins were significantly enriched compared to the control as shown in the Volcano plot (**Fig. 1b**). From all three pulldown experiments, we found 28 proteins significantly enriched in at least two replicates and 8 proteins enriched in all three replicates (**Fig. 1c and Table 1**). Amongst these proteins, several are considered conceivable cellular binding partners based on their cellular localization or function (**Table 1**). Several are known to be palmitoylated and might therefore act as substrates of DHHC20. For example, Linker for activation of T cells (LAT) is a well described palmitoylated protein that was robustly found enriched in our experiment^19^. Also, Phosphatidylinositol 4-kinase type 2-alpha (PI4K2A), a soluble protein that becomes membrane-resident upon palmitoylation, was identified^20^. In addition, subunits of the family of G proteins were significantly enriched, indicating their importance as signaling molecules that require reversible membrane recruitment to initiate signaling events^21^. Notably, the most highly enriched protein is the Transmembrane Protein 109 (TMEM109), devoid of cysteine residues, rendering it non-susceptible to DHHC20 palmitoylation. The protein is a monoatomic cation channel permeable to potassium and calcium^22^, with unknown function in T cells. Its strong enrichment argues for it to be a stably bound interactor of DHHC20. In summary, our results indicate that our approach could identify stably bound interacting proteins as well as known and putative substrates of DHHC20.

**Table 1:** List of proteins significantly enriched in DHHC20 pulldown experiments with Jurkat T-cell lysates, identified in at least two out of three independent experiments. Proteins with at least a 4-fold increase in relative intensity compared to control (log2 fold difference >2) and a q-value<0.05 were considered significantly enriched.

| Gene Name | Function | Palmitoylation (reported) |
| --- | --- | --- |
| DHHC20-Strep | Human Palmitoyltransferase tagged with a TwinStrep-Tag for purification<sup>10</sup> | Palmitoylated<sup>10</sup> |
| TMEM109 | Is a subunit of the RNA exosome complex and is proposed to play a role in the ER $\text{Ca}^{2+}$ release<sup>23</sup> | - |
| HSPA1B | Heat shock protein, a molecular chaperone that is expressed in response to stress<sup>24</sup> | Palmitoylated<sup>24</sup> |
| ICAM2 | Tumor suppressor protein that regulates lymphocyte recirculation<sup>25</sup> | - |
| PI4K2A | Functions as a key regulator of lipid transfer in ER-membranes<sup>26</sup> | Palmitoylated<sup>26</sup> |
| DNAJC13 | Functions in membrane trafficking through early endosomes<sup>27</sup> | - |
| RALA | Ras-like small GTPase<sup>28</sup> | Palmitoylated<sup>28</sup> |
| PGRMC1 | May act as a chaperone and shuttles newly synthesized heme from mitochondria to cytochrome P450 enzymes<sup>29</sup> | - |
| TMX1 | Protein disulphide isomerase family member, palmitoylation enables its trafficking to mitochondria-associated membrane<sup>30</sup> | Palmitoylated<sup>30</sup> |
| LAT | Signaling molecule in T cell activation<sup>31</sup> | Palmitoylated<sup>31</sup> |
| FAM162A | Proposed to be involved in regulation of apoptosis<sup>32</sup> | Palmitoylation predicted<sup>32</sup> |
| CD81 | CD81 belongs to the tetraspanin superfamily and is heavily palmitoylated in the juxtamembrane cysteine residues<sup>33</sup> | Palmitoylated<sup>33</sup> |
| ATL3 | ER- and Golgi-resident GTPase atlastin-3, its co-factor is known to be palmitoylated<sup>34</sup> |  |
| NDC1 | A nucleoporin that functions at the nuclear pore complex<sup>35</sup> | Palmitoylation predicted<sup>36</sup> |
| TMEM43 | In complex it is able to modulate the innate immune signalling through the cGAS-STING pathway<sup>37</sup> | - |
| THEM6 | A ER membrane-associated protein that regulates intracellular levels of ether lipids and plays a key role in the induction of the ER stress response<sup>38</sup> | - |
| APMAP | A novel regulator of extracellular matrix components that interacts with lysyl oxidase-like 1 and 3<sup>39</sup> | - |
| SGPL1 | Cleaves sphingosine-1-phosphate into fatty aldehydes and phosphoethanolamine. Elevates stress-induced ceramide production and apoptosis<sup>40,41</sup> | - |
| MOGS | First enzyme in the N-linked oligosaccharide processing pathway, cleaves the distal alpha 1,2-linked glucose residue from the Glc3-Man9-GlcNAc2 oligosaccharide precursor highly specifically<sup>42</sup> | - |
| RAB14 | A Rab GTPase that plays a role in Glut4 membrane trafficking<sup>43,44</sup> | Palmitoylated<sup>43</sup> |
| GNB2 | Involved as a modulator or transducer in various transmembrane signaling systems. Required for the GTPase activity, for replacement of GDP by GTP, and for G protein-effector interaction<sup>45</sup> | - |
| DDOST | Encodes a component of the oligosaccharide transferase complex and is related to the N-glycosylation of proteins<sup>46</sup> |  |
| MLEC | Classified as a chaperone and a carbohydrate-binding protein with selectivity for the Glc2-N-glycan ER intermediate<sup>47,48</sup> | Palmitoylated<sup>47</sup> |
| PREB | Transcription factor involved in gene regulation, RNA processing, cytoskeleton assembly, vesicle trafficking, and signal transduction<sup>49-51</sup> | - |
| ATAD3B | May act as adaptor of mitochondrial and nucleoid cholesterol homeostasis and metabolism in hESCs and cancer cells<sup>52</sup> | - |
| GNB1 | Involved as a modulator or transducer in various transmembrane signaling systems. Required for the GTPase activity, for replacement of GDP by GTP, and for G protein-effector interaction<sup>45</sup> | - |
| RAB5C | A small GTPase that is a key component of the molecular machinery which is able to drive CD93 recycling to the endothelial cell surface<sup>53</sup> | Palmitoylated<sup>54</sup> |
| RAB1B | A small G-protein Rab1b, which acts as a central regulatory hub in vesicular trafficking<sup>55</sup> | Palmitoylated<sup>56</sup> |

**Figure 1:**
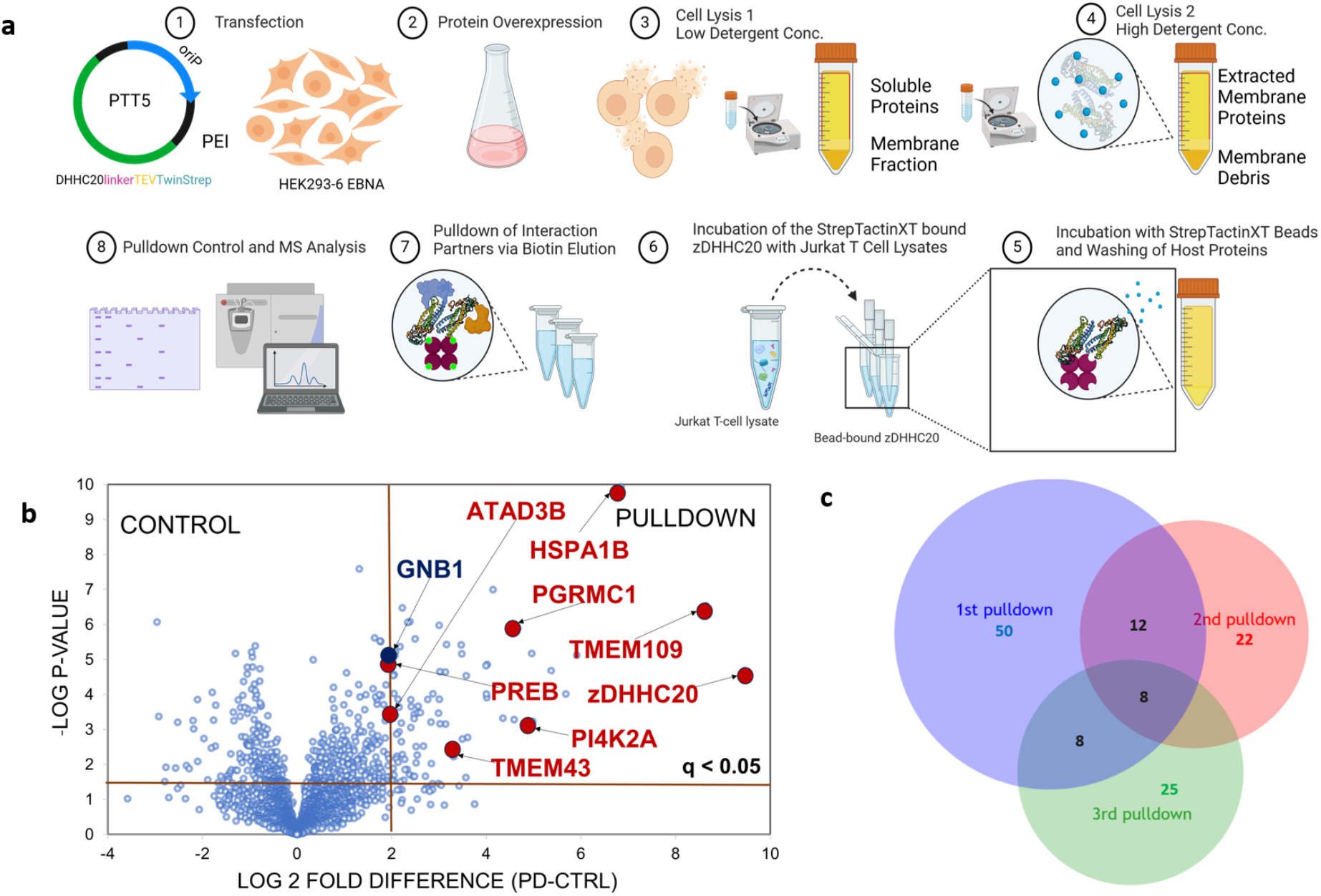
Enrichment of DHHC20 and pulldown experiments. **a**. Flowchart illustrating the experimental procedure for DHHC20-Strep overexpression in HEK293-6 EBNA cells, subsequent purification and immobilization utilizing StrepTactinXT beads, and the pulldown experiment involving the interaction of bead-bound DHHC20-Strep with Jurkat T-cell lysates to identify interaction partners. **b**. Volcano plot of pulldown experiment 1 showing the relative protein intensity difference of proteins enriched by Strep-tagged DHHC20 (Pulldown) compared to the empty bead control (Control). Significant hits are shown in the top right corner indicated by the brown lines (q-value<0.05 and log_2_ fold difference >2). Proteins, given as gene names, that were enriched in all three independent pulldown experiments are highlighted in red. The beta subunit of the heterotrimeric G protein Gαsβ1γ2, is depicted in dark blue, since it appeared in two out of three pulldowns. Entire protein lists of all three pulldown experiments are shown in **Table S1a-c. c**. Venn diagram illustrating the overlap of significantly enriched proteins identified from three independent pulldown experiments. A total of 8 proteins were found to be consistently enriched across all three experiments, while 28 proteins were enriched in at least 2 out of 3 experiments (**Table 1**).

### Cross-linking mass spectrometry of DHHC20 and a trimeric G protein

Two of the enriched proteins from the pulldown experiments were Gβ1 and Gβ2, the beta subunits of the heterotrimeric G proteins Gαsβ1γ2 and Gαsβ2γ2. Trimeric G proteins are known to be palmitoylated at an N-terminal cysteine residue of the α subunit and they were previously found as a class of proteins that are inducibly palmitoylated upon T cell stimulation^57^. We thus decided to investigate the direct interaction between DHHC20 and Gαsβ1γ2 in more detail by a cross-linking experiment combined with MS (XL-MS). For complex formation, recombinantly expressed and purified proteins were employed (**Fig. S1**).

The MS-cleavable cross-linker Disuccinimidyl Dibutyric Urea (DSBU) was utilized, which contains two reactive groups that react specifically with primary amines and therefore covalently link lysine side chains that are in close proximity^58^. The maximum C_α_-C_α_ distance of lysine residues cross-linked by DSBU is estimated to be around 30 Å which results from the fixed length of DSBU (12.5 Å) plus the length of two lysine side chains (12.8 Å) and an estimated tolerance between 2 and 5Å, thus providing soft distance constraints^59^. After the cross-linking reaction, proteins were digested into peptides using trypsin and crosslinked peptides from two independent experiments were identified by LC-MS (**Fig. 2a and Table S2**).

**Figure 2.**
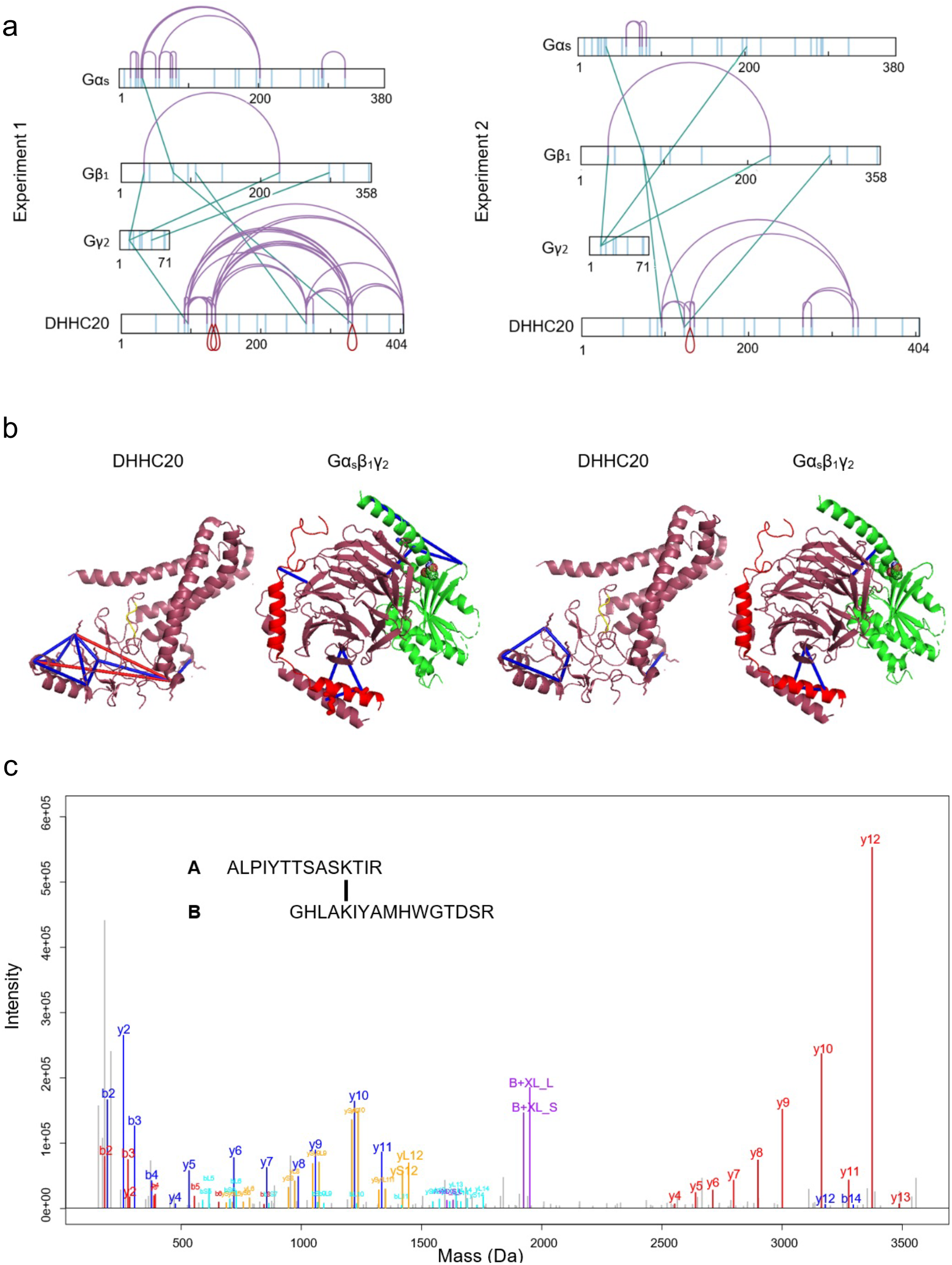
Cross-linking of DHHC20 with the heterotrimeric G protein Gαsβ1γ2. **a**. Cross-links detected by MS between DHHC20 and Gαsβ1γ2 from two independent experiments. The vertical lines represent the distribution of lysine residues within the protein sequence. Interprotein cross-links are indicated by straight lines, while self-links (inter- or intra-molecular) are indicated by curved lines. The image was generated with xiNet^60^. The numbering of Gβ1 in the Figure is shifted by 18 additional amino acids due do an N-terminal expression tag (compared to the sequence from UniProt-ID: P54311). **b**. Illustration of the intra cross-links within DHHC20 (PDB: 6BML) and Gαsβ1γ2 (PDB: 7PIU) from two experiments, validating their structural integrity. Bromopalmitate was removed from the DHHC20 structure, and the melanocortin-4 receptor was omitted from the G protein model, aligning with the absence of these entities within the experimental framework of our investigation. The subunits of the G protein are depicted in the figure according to the following color code: alpha in green, beta in raspberry and gamma in red. Blue lines denote intraprotein cross-links within the 30 Å cross-linker restraint, while red lines exceed this limit. **c**. Example MS2 spectrum confirming the cross-link formed between Lys_123_ of DHHC20 (peptide A) and Lys_280_ of Gβ1 (peptide B). All peaks are charge-deconvoluted. Characteristic b- and y-fragment ions resulting from cleavage of peptide A and B are labeled in red and blue, respectively. Signature peaks of peptides A and B resulting from the cleavage of the DSBU cross-linker are labeled in purple (S/L indicates whether the peptide contains the shorter/longer arm of DSBU after cleavage). Characteristic b- and y-fragment ions resulting from backbone cleavage of peptide A or peptide B and additional cleavage of the DSBU cross-linker are labeled in orange and cyan, respectively.

Results from two independent biological replicates are shown in **Fig. 2a**, displaying (i) intraprotein cross-links within DHHC20, (ii) intraprotein cross-links within individual G protein domains, (iii) intersubunit cross-links between the α, β and γ domains of the G protein and (iv) interprotein cross-links between DHHC20 and individual domains of the G protein. The intraprotein cross-links within DHHC20 or between G protein domains served to probe the specificity of cross-links and at the same validate the structural integrity of the proteins by comparison with existing structures. For DHHC20 an X-ray structure was used for comparison (PDB: 6BML)^10^, while for Gαsβ1γ2 (PDB: 7PIU), an electron microscopy structure of the protein in complex with the melanocortin-4 receptor was chosen as a reference (**Fig. 2b**). Among the 11 intraprotein cross-links detected within the structured region of DHHC20 across the two experiments, only two violated the distance restraint set by the cross-linker in the initial experiment. We assume that these violations may be due to conformational flexibility of the region containing the respective lysines or the formation of DHHC20 homodimers. For the trimeric G protein, 9 cross-links between subunits were identified in total, with all of them obeying the cross-link distance constraints. Thus, the cross-links reproduce the known structures of DHHC20 and the trimeric G protein.

In total, six interprotein lysine cross-links involving DHHC20 were detected from the two experiments. Five cross-links were observed between DHHC20 and Gβ1 (Lys_96_-Lys_57_, Lys_123_-Lys_57_, Lys_123_-Lys_280_, Lys_265_-Lys_89_, Lys_331_-Lys_57_), while one cross-link was detected between DHHC20 and Gg_2_ (Lys_96_-Lys_14_). The charge-deconvoluted annotated MS2 spectra from all interprotein lysine cross-links are shown in **Fig. 2c and Fig. S2**.

These diagnostic cross-links were used to perform rigid docking by HADDOCK^61^ with the cross-links as soft constraints. From the two sets of cross-links two clusters of possible orientations were obtained and then energy minimized. Representative structures of the docking poses from the two experiments are shown in **Fig. 3** and corroborate the β-barrel of Gβ as the main site of interaction. From the two sets of cross-links, two sets of possible orientations were obtained and then energy minimized (see methods section for detail). Representative structures of the docking poses are shown in **Fig. 3**. The two most likely poses consistent with the cross-links from experiment 1 (**Fig 3A and 3B**) are similar to each other, evident from their superposition (**Fig. 3C**). By using the cross-links from experiment 2 as constraints, the two most likely poses (**Fig. 3D and 3E**) are quite different from each other (**Fig. 3F**) and from the two poses from experiment 1 (**Fig. 3G**).

**Figure 3.**
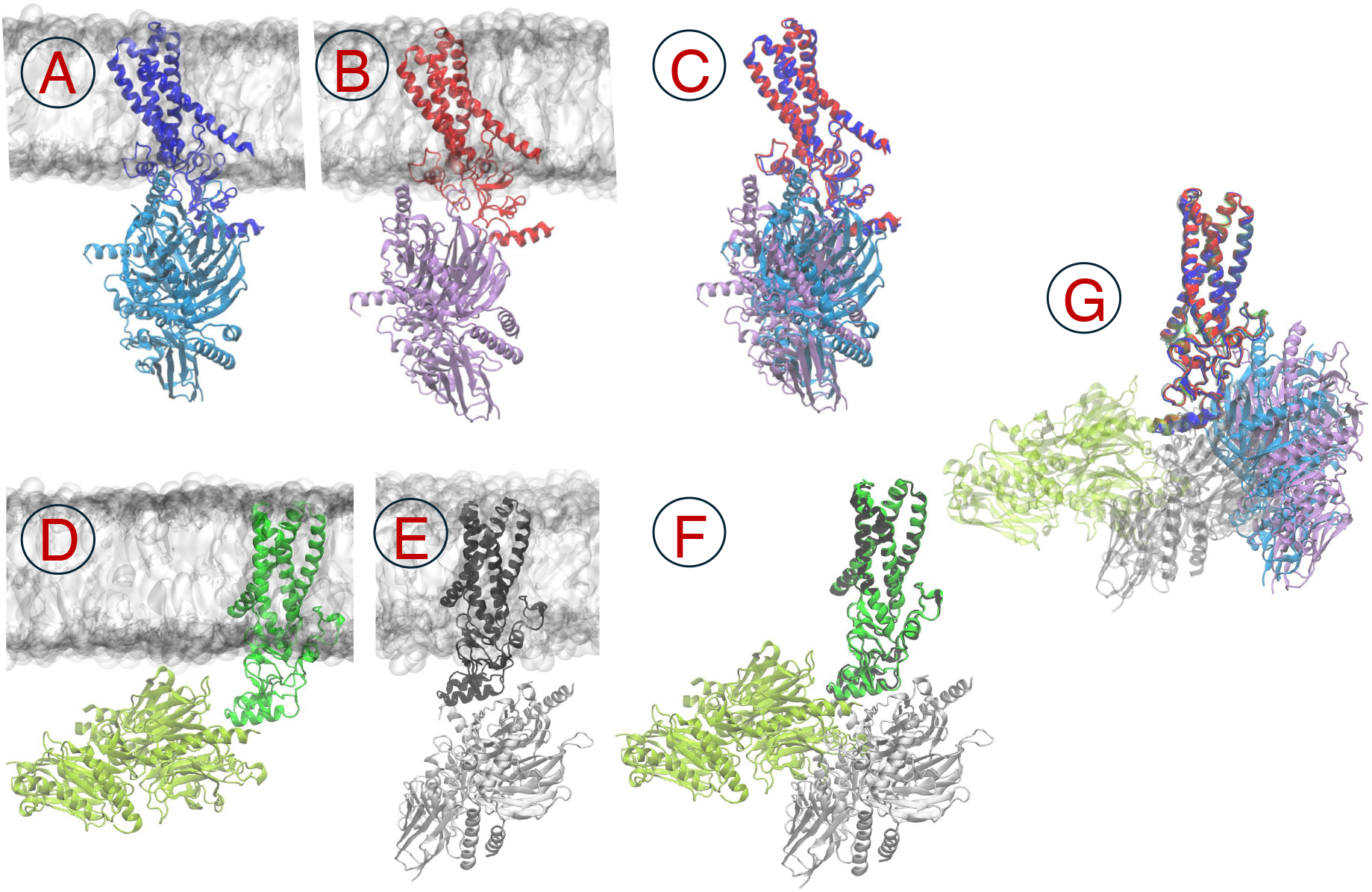
Results from docking simulations. **A** and **B** show the two most likely poses resulting from docking the two proteins imposing the Xlinks from experiment 1 as constraints. In **A**, DHHC is colored in blue and G protein in cyan, while in **B** DHHC is colored in red and G protein in purple. **C** shows the superposition of pose **A** and **B**. The membrane is omitted for clarity. **D** and **E** show the two most likely poses resulting from docking the two proteins imposing the Xlinks from experiment 2 as constraints. In **D** DHHC is colored in bright green and G protein in lime green, while in **E** DHHC is shown in black and G protein in grey. **F** shows the superposition of poses **D** and **E**. Panel **G** shows all four poses superimposed.

## Discussion

Defining the substrate specificity of DHHC enzymes is a major challenge in the field of catalyzed palmitoylation^62^. The principal architecture of this family of proteins has been elucidated in a seminal paper by the Banerjee group and the placement of the transferable palmityl chain was identified by localizing the irreversible inhibitor 2-bromo-palmitate in the active site^10^. While it is clear that the lipid moiety has to be transferred from the catalytic cysteine to a cysteine of the incoming substrate, there is no structural information of how substrates are steered towards the active site. No clear rules have emerged based on sequence comparison of palmitoylation sites, highlighting the importance of obtaining structural information. Considering the structural diversity of substrates, encompassing palmitoylation sites residing in both structured and flexible domains, spanning transmembrane or soluble proteins, and occurring at various cellular locations such as the endoplasmic reticulum (ER), Golgi, or plasma membrane, addressing this challenge requires the examination of numerous enzyme-substrate complexes to conceptualize the mode of action. Furthermore, proteins interacting with DHHC proteins may not serve as substrates but rather stabilize the enzyme or modulate substrate specificity by contributing to substrate recognition^13,14,63^. Thus, the challenge is on the one hand to identify interacting proteins and then to map the molecular interface of either stably binding interaction partners or substrates.

Here, we took a step towards these goals by performing pulldown experiments with purified DHHC20 to identify putative interaction partners, subsequently confirming a specific interacting protein by mapping structural interfaces through cross-linking mass spectrometry. The list of interactors contained transmembrane as well as peripheral membrane proteins. Interestingly, the protein with the highest enrichment factor was the cation channel TMEM109 (mitsugumin-23)^22^ localized to organellar membranes and transporting potassium as well as calcium ions. TMEM109 does not contain a single cysteine, thus it is certainly not a target of DHHC20. It therefore represents a candidate for a DHHC20 chaperone and suggests a further link between DHHC20 and calcium signaling^17^. High enrichment factors were further on observed for the single-pass transmembrane proteins ICAM-2 and the progesterone receptor PGRMC1. Both feature an LC(I,F)(F,I) motif at the C-terminal part of their transmembrane helix in proximity to the cytoplasm. It would be interesting to investigate whether these cysteines can be palmitoylated and whether DHHC20 serves as a palmitoyl transferase in these instances. Of the soluble interaction partners that were enriched at least two out of three times in our experiments several are known to be palmitoylated. For example, the PI4 kinase 2A contains a CCPCC motif that was shown to be palmitoylated by DHHC3 and DHHC7 in the trans-Golgi^23,24^. Co-immunoprecipitation depended on palmitoylation of the motif and on cholesterol. We therefore propose that in T cells DHHC20 may serve as the interaction partner and putative PAT for PI4K2A. Another protein known to be palmitoylated is the family of Gα subunits of trimeric G proteins, a protein family that was also shown to be inducibly palmitoylated upon T cell stimulation^19^. The N-terminus of the Gα domain features a cysteine known to be palmitoylated^64^, for example by DHHC5 in cardiomyocytes^65^ and is of importance for downstream functions as for example β-adrenergic signaling. In contrast, the β1 and γ2 subunits of the G protein are not palmitoylated but reside at the membrane by geranyl-geranylation of a cysteine of the CAAX box in the γ2 subunit^21^. In our pulldown experiment we found the Gβ1 subunit of Gαs-short in two of the experiments and the Gγ2 subunit in one of the experiments. We therefore conclude that it is the the βγ heterodimer that is stably associated with DHHC20 in T cells. Interestingly, previous studies have suggested that DHHC3 and DHHC7 can palmitoylate Gα proteins in HEK293 cells^66^, while DHHC5 is suggested to be the PAT for Gαs and Gαi in cardiomyocytes^65^. It will be very interesting to see whether palmitoylation by these enzymes may also occur through interaction of the βγ subunits of the trimeric G protein.

To further test the hypothesis of a direct interaction, we used cross-linking mass spectrometry as a method to probe whether the DHHC20 protein and the trimeric G protein are approaching each other at a distance smaller than approximately 30 Å, as defined by the chemical structure of the lysine-cross-linker DSBU. Observing intra-protein lysine cross-links that are largely within the length limit gave confidence into data interpretation, on the one hand by corroborating the intactness of the fold of the two proteins and on the other hand showing that unspecific cross-links are the exception. In two independent experiments it was the β1-subunit that resulted in either 2/3 or 3/3 interprotein cross-links to the DHHC20 enzyme, in agreement with the β1 subunit to be the most highly enriched domain found in the pulldown experiments. Interestingly, the two sets of interprotein cross-links were different in the two experiments, but both involved residue K96 of DHHC20 and K57 of Gβ as common denominators of the interface. Two docking clusters can be visualized that probably represent possible orientations of how the two proteins interact. It will now be necessary to challenge these models by high resolution structural methods such as X-ray crystallography or electron microscopy. Nevertheless, given the routine application of XL-MS in deriving valid restraints for structural modeling or verification of known structures^67,68^, we think that the methodology can be used in a meaningful way to help decipher the rules of substrate recognition by DHHC enzymes in the future.

## Methods

### Expression and purification of DHHC20

The HEK293 cell line stably expressing the Epstein-Barr virus nuclear antigen-1 (HEK293-EBNA1, or 293-6E) was used for large-scale transfection. The construct of interest (DHHC20-linker-TEV-TwinStrep) was cloned in a pTT expression vector, which bears the Epstein-Barr virus origin of replication, oriP, leading to a threefold increased protein yield compared to a non-oriP vector^69^. HEK293-6 EBNA cells were passaged in HEK TF medium from Xell AG, supplemented with 6 mM L-Glutamine, and expanded until a concentration of 2-3×10^6^ cells/mL and a viability of 100% was reached. The cells were pelleted down and resolved in HEK TF medium to reach a concentration of 5×10^6^ cells/mL. A DNA/PEI mix was added and after 4h of shaking at 120 rpm at 37°C, 5% CO_2_, medium was added to dilute the cells to 2.5×10^6^ cells/mL. For the transfection, the DNA/PEI mix was prepared in 10% volume of the final transfection volume. The DNA/PEI ratio required was 1 μg DNA and 2 μg PEI per million cells, referring to the first step of transfection before dilution. The cells were left to grow for two days, after which a maximal expression level of protein was expected (determined by ELISA). The cells were then pelleted down, the pellet weight was determined, and the pellet was snap frozen in liquid nitrogen and stored at -80°C until further use.

The transfected HEK cell pellet was allowed to thaw on ice and subsequently lysed using an initial low detergent concentration lysis buffer (300 mM NaCl, 0.1% DDM, protease inhibitor (1:100), 100 mM TRIS pH 8, 5 mM DTT) aiming to extract all soluble proteins. Approximately 6 mL of lysis buffer per gram of cells was employed. The lysis mixture underwent a 1-hour incubation at 4°C with rotation, followed by a 30-minute centrifugation at 4°C, 10,000 rcf using a SA 25.5 rotor. The resulting supernatant was discarded, and the pellet underwent a second lysis using a new buffer characterized by an elevated detergent concentration (300 mM NaCl, 1% DDM, protease inhibitor (1:100), 100 mM TRIS pH 8, and 5 mM DTT). This second lysis involved thorough vortexing of the pellet and a 1-hour incubation at 4°C with rotation. Subsequent to this, another centrifugation at 4°C, 10,000 rcf in a SA 25.5 rotor ensued. Meanwhile, 100 μL of high-capacity Strep-TactinXT 4Flow beads (in 50% suspension) from Iba-Lifesciences were centrifuged at room temperature for 5 minutes at 1000 rcf and washed once with 1 mL washing buffer (300 mM NaCl, 0.05% DDM, 100 mM TRIS pH 8, and 5 mM DTT). Following centrifugation, the cleared lysate was incubated with the pre-washed beads for 1.5 hours at 4°C with rotation. Subsequently, the mixture of beads and cleared lysate underwent a 5-minute centrifugation at 4°C, 1000 rcf. The resulting supernatant was subjected to a second incubation with 100 μL freshly washed beads, followed by a 1.5-hour incubation at 4°C with rotation and a subsequent 5-minute centrifugation at 4°C, 1000xg.

Post-centrifugation, each batch of beads underwent three washes with 1 mL washing buffer, with the supernatant being discarded after each wash. In the last step, 800 μL washing buffer was added to the resulting 200 μL of beads and 50 μL (corresponds to 10 μL settles beads) suspension was taken out and transferred to a new tube. The suspension was centrifuged for 5 mins at 1000 rcf and the settled beads were eluted with 30 μL elution buffer. 20 μL eluate was mixed with 5 μL sample buffer with ß-mercaptoethanol and boiled for 5 min. An SDS gel was then run as a control for the protein expression. The elution of the rest of the beads was accomplished using three times the volume of the beads of elution buffer (100 mM TRIS pH 8, 150 mM NaCl, 50 mM biotin, 0.05% DDM). The resulting supernatant, containing the purified protein, was carefully transferred to a fresh Eppendorf tube, while the beads were regenerated in MgCl_2_ buffer for subsequent use.

### Expression and purification of Gαsβ1γ2

For the production of the heterotrimeric G protein Gαsβ1γ2, previously published constructs were used^70^. For this purpose, the sequence of the bovine Ga_s_ short subunit (UniProt-ID: P04896-2) was cloned into the backbone of pFastBac vector and rat Gβ1 (UniProt-ID: P54311, supplemented with a Hexa-His-tag) and bovine Gγ2 (UniProt-ID: P63212) subunits were inserted into the pFastbacDual vector. Virus generation was performed using the Bac-to-Bac™ expression system (Thermo Fisher Scientific) in *Sf*9 insect cells. After extensive titrations to determine the optimal virus ratio, the heterotrimeric G protein was expressed after co-infection with Gαs and Gβ1γ2 viruses in *Trichoplusia ni* (Tni) insect cells for 40 h at 28°C^71^. Following cell harvest by centrifugation, the cell pellet was stored at -20 °C until use. The purification was mainly based on the protocol published by Rasmussen and colleagues^70^, which started with the addition of lysis buffer (10 mM TRIS pH 7.5, 0.1 mM MgCl_2_, 2 μM leupeptin, 1 mM benzamidine, 0.5 mM PMSF, 5 mM 2-mercaptoethanol and 10 μM GDP) to the cell pellets. After incubation for 30 min the cell suspension was centrifuged for 30 min at 50,000 rcf and the pellet was resuspended in solubilization buffer (containing 0.05% DDM/CHS, 1% Na-cholate and 15% glycerol) and solubilized for 1 h at 4 °C. After renewed centrifugation, the heterotrimeric G proteins were bound to Ni-NTA beads for 1 h at 4 °C. Unspecific bound proteins were washed away by several washing steps with increasing concentrations of imidazole and NaCl before elution with a gradient up to 180 mM imidazole. The fractions containing G protein (determined by SDS-PAGE) were pooled, concentrated, and incubated with λ-phosphatase, CIP and antarctic phosphatase. This was followed by a final purification step using a MonoQ column. After elution with high salt buffer, the purest fractions were combined, pooled, snap frozen in liquid nitrogen and stored at -80 °C until use.

### Jurkat T-cell culture and lysis

A 4 million-cell containing aliquot of Jurkat T-cell E6 clone was thawed and cultured in RPMI medium (Biochrom), supplemented with L-glutamine and 10% fetal calf serum. The Jurkat T-cell stock was maintained at a concentration ranging between 0.2 and 1 million cells/mL. For expansion, T300 bottles were utilized, each containing 100 mL of Jurkat T-cell stock with an average concentration of 2.5 million cells/mL. The expanded cells were pelleted by centrifugation at 300 rcf, RT, for 7 minutes. The supernatants were carefully removed, and the cells were subjected to two washes with PBS. Subsequently, aliquots of 500 million cells each were shock-frozen in liquid nitrogen and stored at -80°C. On the day of the experiment, the aliquots were allowed to thaw at room temperature. Each thawed aliquot was lysed in 500 μL of lysis buffer (50 mM NaH_2_PO_4_ pH 7.2, 150 mM NaCl, 1% DDM, protease inhibitor 1:100, 1 mM PMSF, DNase, RNase). The lysate was incubated on ice for 30 minutes, followed by the clearing of lysates through several centrifugation steps at maximum speed for 5 minutes at 4°C.

### Pulldown of DHHC20 with Jurkat T-cell lysates

In the context of this experiment, the purification of DHHC20, as detailed above, was carried out until the point where the protein was immobilized on Strep-Tactin XT agarose beads. The protein-bound beads were dispensed into four aliquots, each containing 45 μL of beads. Additionally, four aliquots comprising 45 μL of unbound beads were utilized as control samples to assess background protein levels in the pulldown experiment. Subsequently, the DHHC20-bound and control beads were subjected to two washes with 500 μL of Jurkat T-cell lysis buffer (50 mM NaH_2_PO_4_ pH 7.2, 150 mM NaCl, 1% DDM, protease inhibitor 1:100, 1 mM PMSF, DNase, RNase). This step aimed to ensure uniformity in buffer composition between the beads and the Jurkat T-cell lysates for the subsequent pulldown procedure. For each pulldown, 45 μL of DHHC20-bound and control beads were combined with 500 μL of Jurkat T-cell lysate. All pulldown samples were then subjected to an overnight incubation at 4°C on an end-over-end shaker to facilitate the formation of complexes between DHHC20 and potential binding partners from the T-cell lysate. The protein complexes obtained from the overnight incubation were subjected to two washes using DHHC20 washing buffer (300 mM NaCl, 0.05% DDM, 100 mM TRIS pH 8, 5 mM DTT) and subsequently eluted with 135 μL elution buffer (100 mM TRIS pH 8, 150 mM NaCl, 50 mM biotin).

The samples were then prepared for mass spectrometry measurements. To facilitate denaturation and reduction of disulfide bonds, the proteins were treated with 300 μL 2 M urea containing 5 mM DTT in a 100 mM ammonium bicarbonate buffer for 30 minutes at 56°C. Subsequently, free thiols were alkylated by adding 30 μL of 100 mM iodoacetamide, resulting in a final concentration of 8 mM iodoacetamide. The alkylation reaction was incubated at room temperature for 30 minutes in the dark. The digestion was performed by adding 0.2 μg Trypsin with overnight incubation at 37°C. After digestion, salts and detergents were removed before LC-MS analysis by SDB-RPS StageTips (Empore™ 2241) as described previously^72^. Eluted peptides were dried by vacuum centrifugation and stored at -20°C.

### Direct cross-linking of DHHC20 with Gαsβ1γ2

DHHC20 and Gαsβ1γ2 were mixed at equimolar concentrations (6 μM each) in a final reaction volume of 70 μL. The proteins were cross-linked at room temperature by adding 0.7 μL Disuccinimidyl Dibutyric Urea (DSBU) (from a 50 mM stock solution in DMSO) twice, in two sequential steps, each lasting 20 minutes. Subsequently, the reaction was quenched for 30 minutes at RT by adding 1.45 μL 1M TRIS per sample achieving a final concentration of 20 mM TRIS. Following quenching, 72,85 μL of a 100 mM ABC (ammonium bicarbonate) buffer was added. The proteins were then denatured by adding 70 mg crystalline urea (8 M final urea concentration). Disulfide bonds were reduced by the addition of 0,4 μL 2M DTT (reaching a final concentration of 5 mM DTT) in 100 mM ABC and incubation for 30 minutes at 56°C. Free thiols were subsequently alkylated with 8 mM iodoacetamide (13.7 μL per sample from a 100 mM stock) at room temperature for 30 minutes in the dark. The pH of the mixture was verified to be around 8 before initiating protein digestion with Lys-C at a 1:75 enzyme-to-substrate ratio (0,73 μg Lys-C for 55 μg protein), conducted for 4 hours at 37°C. Following this, the urea concentration was reduced to 2 M by adding 563 μL 100 mM ABC buffer, and the samples were subjected to additional digestion with trypsin at a 1:100 enzyme-to-substrate ratio (0,55 μg Trypsin for 55 μg protein) overnight at 37°C. After digestion, salts and detergents were removed before LC-MS analysis by SDB-RPS StageTips (Empore™ 2241) as described previously^25^. Eluted peptides were dried by vacuum centrifugation and stored at -20°C.

### Nano liquid chromatography-mass spectrometry (LC-MS)

Dried peptides were reconstituted in 20 μL of 0.05% trifluoroacetic acid (TFA), 4% acetonitrile in water, and 5 μL were analyzed by an Ultimate 3000 reversed-phase capillary nano liquid chromatography system connected to a Q Exactive HF mass spectrometer (Thermo Fisher Scientific). Samples were injected and concentrated on a trap column (PepMap100 C18, 3 μm, 100 Å, 75 μm i.d. x 2 cm, Thermo Fisher Scientific) equilibrated with 0.05% TFA in water. After switching the trap column inline, LC separations were performed on a capillary column (Acclaim PepMap100 C18, 2 μm, 100 Å, 75 μm i.d. x 50 cm, Thermo Fisher Scientific) at an eluent flow rate of 300 nL/min. Mobile phase A contained 0.1% formic acid in water, and mobile phase B contained 0.1% formic acid in 80% acetonitrile / 20% water. The column was pre-equilibrated with 5% mobile phase B followed by an increase of 5–44% mobile phase B in 70 min. Mass spectra were acquired in data-dependent mode with the following settings: MS scan range m/z 350–1650; MS resolution 60,000; MS AGC target 3×106; MS/MS scans of the TOP15 precursor ions; MS2 resolution 15,000; MS2 AGC target 1×105; collision energy 27; minimum AGC target 1×103; charge exclusion 1 and >5; dynamic exclusion 20 s. For the analysis of the cross-linked peptides, a longer gradient was used with an increase of 5–44 % mobile phase B in 130 min. Mass spectra were acquired in data-dependent mode with the following settings: MS scan range m/z 375–1575; MS resolution 120,000; MS AGC target 3×106; MS/MS scans of the TOP10 precursor ions; MS2 resolution 60,000; MS2 AGC target 1×105; stepped collision energy 21, 27, 33; minimum AGC target 1×104; charge exclusion 1-3 and >8; dynamic exclusion 30 s.

### MS data analysis

For the pulldown experiments, MS and MS/MS raw data were analyzed using the MaxQuant software package (version 2.0.3.0) with Andromeda peptide search engine^73^. Data were searched against the human reference proteome downloaded from Uniprot on April 8, 2023 (Proteome ID UP000005640 containing 81,791 proteins) using the default parameters except for enabling the options label-free *quantification (LFQ) and match between runs*. Filtering and statistical analysis was carried out using the software Perseus version 1.6.14^74^. Only proteins which were identified with LFQ intensity values in at least 3 out of 4 replicates within at least one of the 2 experimental groups (DHHC20 pulldown or control) were used for downstream analysis. Missing values were replaced from normal distribution (imputation) using the default settings (width 0.3, down shift 1.8). Mean log2 fold protein LFQ intensity differences between experimental groups (DHHC20 pulldown or control) were calculated in Perseus using student’s t-tests incorporating a permutation-based False Discovery Rate (FDR) threshold of 0.05. This process ensured a rigorous evaluation of the differences in protein abundance between the DHHC20 pulldown experiments and the control group. Volcano plots were created by plotting the -log_10_ p-value against the mean log_2_ fold protein LFQ intensity differences (DHHC20 pulldown - control).

For the cross-linking experiments, MS and MS/MS raw data were processed using Proteome Discoverer version 2.4 (Thermo Fisher Scientific) to generate MGF files as described before^28^. By using the add-on MS2 Spectrum Processor, all MS2 spectra were deconvoluted to charge state 1. Identification of cross-linked peptides was achieved by using the MGF files as an input for a stand-alone version of XlinkX v2.0 kindly provided by Dr. Fan Liu^75^ The following settings were used: minimum peptide length = 6; maximal peptide length = 35; DSBU cross-linker mass = 196.0848 Da (short arm = 85.0528 Da, long arm = 111.0320 Da); precursor mass tolerance = 10 ppm; fragment mass tolerance = 20 ppm; fixed modification = Cys carbamidomethylation; variable modification = Met oxidation; enzyme = trypsin; missed cleavages = 3; n-score cutoff = 10^−8^. All spectra were searched against a database generated based on the exact sequences of the constructs of DHHC20 and the heterotrimeric G proteins (Gαs, β1, γ2) as described above.

### Computer simulations

Docking simulations between the G-Protein and DHHC were conducted utilizing the Haddock docking software^61^. We used protein chain A of the pdb structure 6BML for DHHC and chains A, B, G and N of PDB entry 7PIU for the G-Protein. Prior to docking, the 7PIU structure was processed using the available pdb-tools^76^, specifically pdb_reres.py and pdb_selchain.py, to prevent redundant residue numbering across multiple chains. The chains R and P were removed, under the assumption that they are not part of the final complex. Two docking experiments were performed in accordance with the two cross-linking experiments, with the inter cross-linking information utilized as unambiguous distance restraint of 15 Å during docking. A total of 9 clusters comprising 67 structures for the first possible pose and 10 clusters comprising 93 structures for the second possible pose were obtained from docking. The top-scoring structures from each of the two best clusters for each experiment was selected, resulting in four structures overall—two for each cross-linking experiment. These structures underwent further refinement through relaxation with all-atom molecular dynamics simulations. Subsequently, we placed the position and orientation of the DHHC-G protein complex with respect to the bilayer with the help of the PPM web server^77^ available through CHARMM-GUI^78^. The resulting structures were embedded into pre-equilibrated POPC bilayers, solvated with TIP3P water and 150 mM NaCl using the CHARMM-GUI membrane builder^79^. After initial energy minimization for 5000 steps with the steepest descent, six rounds of short equilibration runs were performed according to the standard CHARMM-GUI protocol employing the CHARMM36m force field^80^. For simulations, the 2023 cuda version of the GROMACS simulation package was used^81^. All structures are rendered using VMD^82^.

## Acknowledgements

We thank Dr. Jana Sticht for valuable discussions.

## Supplements

**Supplementary figure 1:**
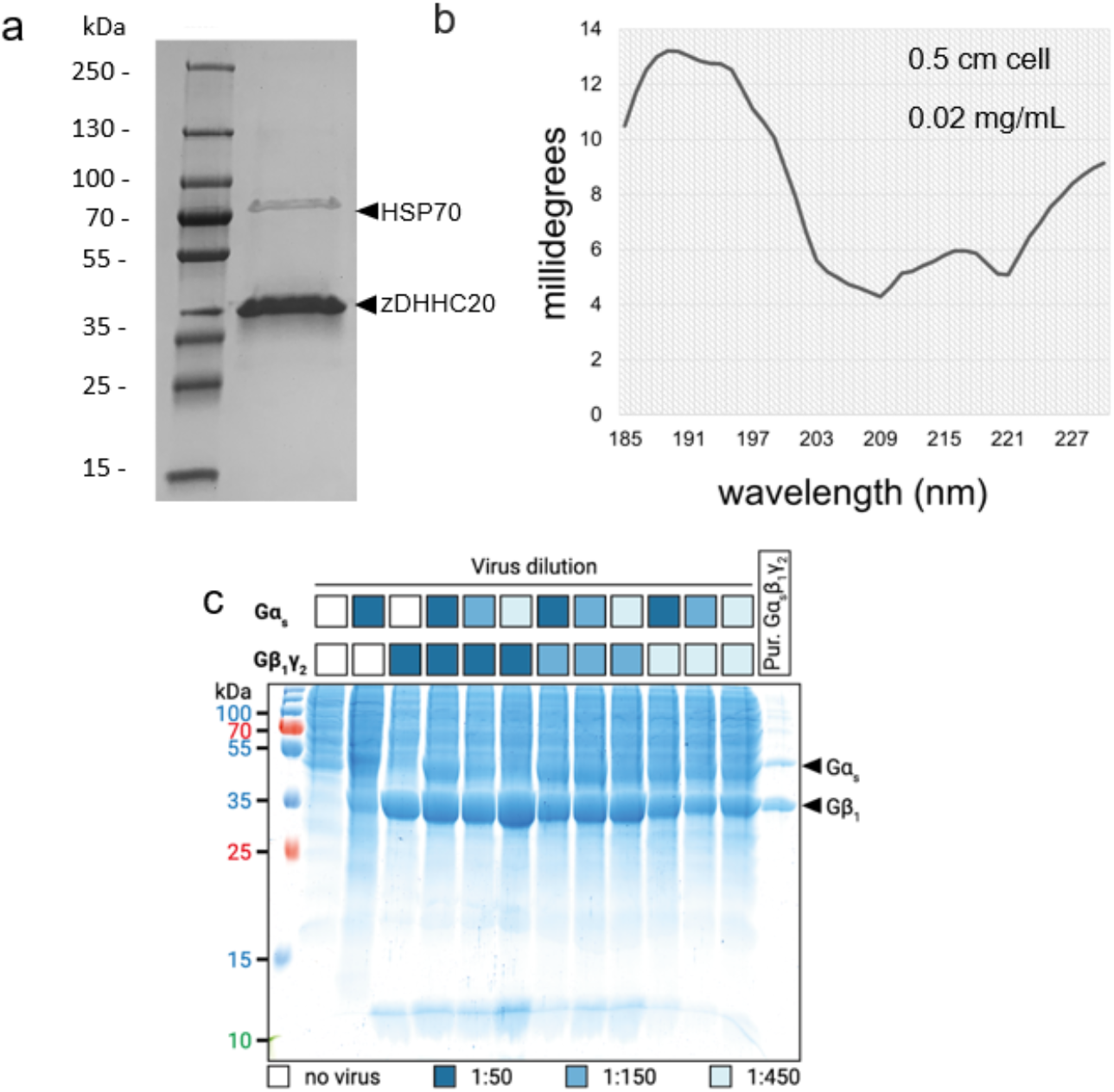
Quality control assessment of recombinantly expressed and purified DHHC20 and Gαsβ1γ2 proteins used for cross-linking MS experiments. **a**. SDS-PAGE of purified DHHC20. Prestained Plus Protein Ladder was used as a molecular weight marker. The gel reveals a well-defined band corresponding to the expected molecular weight of DHHC20 at approximately 46 kDa, confirming the success of the purification process. A second prominent band at around 70 kDa corresponds to Hsp70 (verified by MS). **b**. The CD spectrum of DHHC20 exhibits characteristic features indicative of an alpha-helical protein structure: a minimum at 208 nm, associated with the π → π* transition of the peptide bond and a maximum at 195 nm linked to the n → π* transition of the peptide bond, suggesting a folded and structurally intact protein. **c**. SDS-PAGE of a virus titration experiment and the purified Gαsβ1γ2. Various dilutions of the subunits Gαs and Gβ1γ2 virus were tested to maximize the yield of heterotrimeric G protein. The best ratio was then used for large-scale expression in insect cells, with the last column showing the final sample after completion of an ion-exchange chromatography step. While the Gαs and Gβ1 subunits are clearly detectable, Gγ2 cannot be visualized in the gel due to its small size of 8 kDa.

**Supplementary figure 2:**
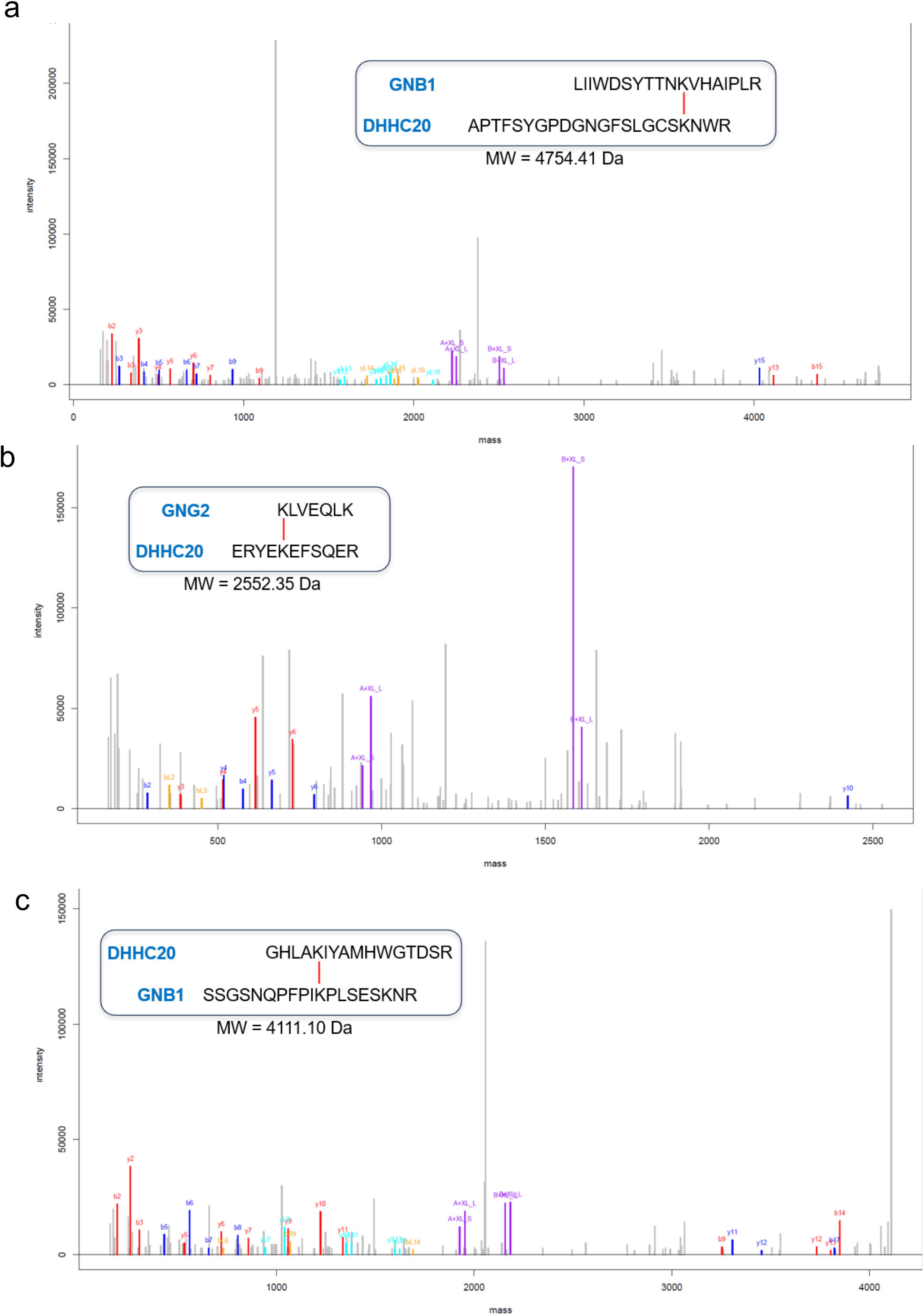
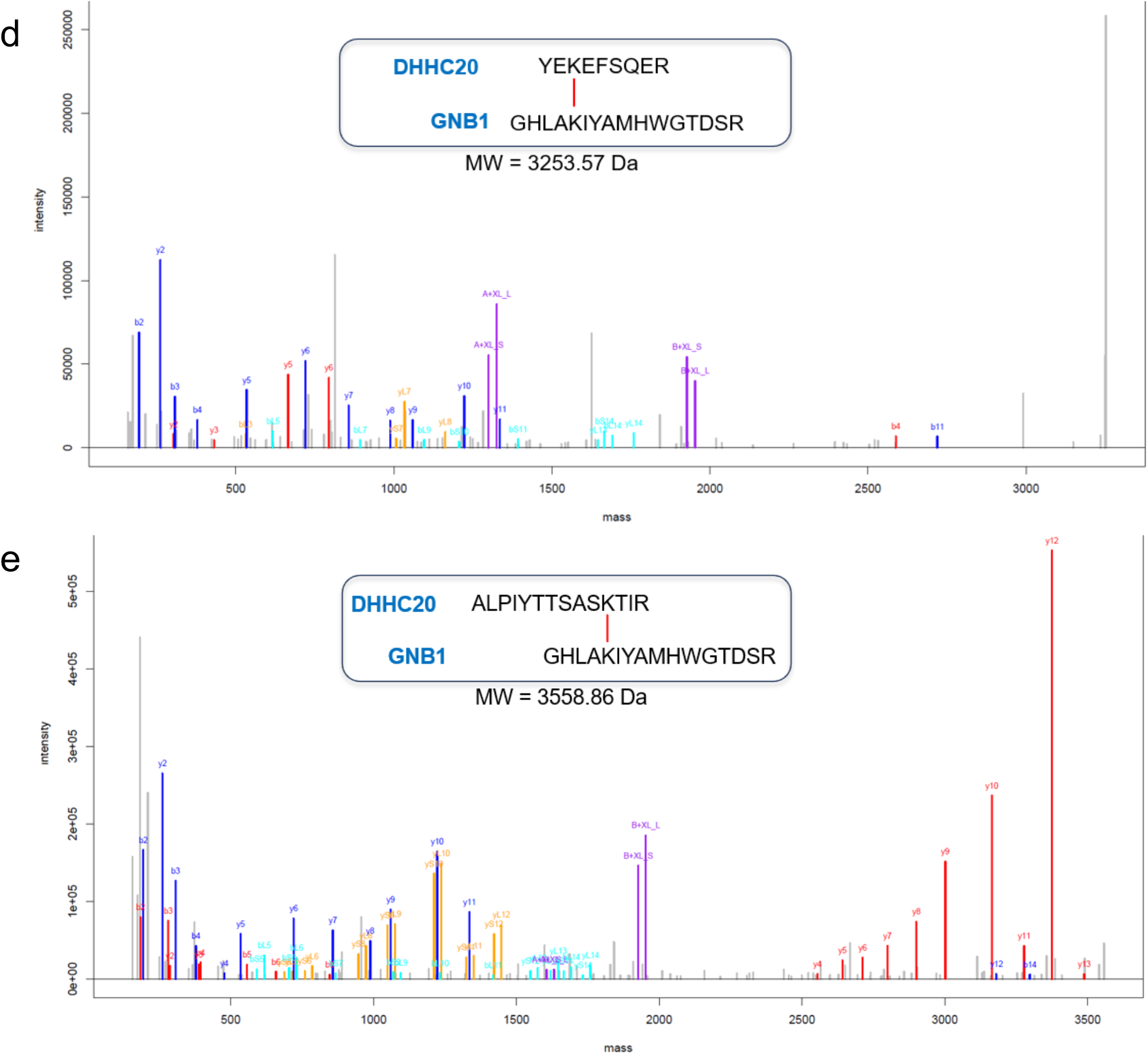
MS2 spectra of cross-linked peptides. Spectra a-c belong to experiment 1 and spectra d and e belong to experiment 2. All peaks are charge-deconvoluted. Characteristic b- and y-fragment ions resulting from cleavage of peptide A (top) and B (bottom) are labeled in red and blue, respectively. Signature peaks of peptides A and B resulting from the cleavage of the DSBU cross-linker are labeled in purple (S/L indicates whether the peptide contains the shorter/longer arm of DSBU). Characteristic b- and y-fragment ions resulting from backbone cleavage of peptide A or peptide B and additional cleavage of the DSBU cross-linker are labeled in orange and cyan, respectively. **a**. MS2 spectrum confirming the cross-link formed between Lys_265_ of DHHC20 and Lys_89_ of Gβ1. **b**. MS2 spectrum confirming the cross-link formed between Lys_96_ of DHHC20 and Lys_14_ of Gγ2. **c**. MS2 spectrum confirming the cross-link formed between Lys_331_ of DHHC20 and Lys_57_ of Gβ1. **d**. MS2 spectrum confirming the cross-link formed between Lys_96_ of DHHC20 and Lys_57_ of Gβ1. **e**. MS2 spectrum confirming the cross-link formed between Lys_123_ of DHHC20 and Lys_57_ of Gβ1

